# A division-associated envelope protein, MAB_2363, drives intrinsic resistance and virulence in *Mycobacterium abscessus*

**DOI:** 10.64898/2025.12.18.695127

**Authors:** Lijie Li, Md Shah Alam, Chunyu Li, H.M. Adnan Hameed, Buhari Yusuf, Aweke Mulu Belachew, Xirong Tian, Abdul Malik, Cuiting Fang, Yanan Ju, Jingran Zhang, Liqiang Feng, Wei Yu, Shuai Wang, Tianyu Zhang

**Author notes:** Correspondence: Shuai Wang, Tianyu Zhang.

## Abstract

*Mycobacterium abscessus* exhibits intrinsic resistance to conventional antibiotics, significantly limiting treatment options. Our previous studies established that MAB_2362 (SteA) is a key regulator of cell division that contributes to intrinsic resistance and virulence. Considering that SteA-like proteins often act alongside SteB counterparts, we hypothesized that the adjacent gene *MAB_2363* encodes the corresponding SteB-like division regulator. In this study, we found that deletion of *MAB_2363* significantly increased susceptibility to multiple antibiotics and disrupted cell wall permeability. Microscopy revealed striking cell division defects in the mutant, including elongated cell morphology and multiple septa. Subcellular localization of a GFP-MAB_2363 fusion protein demonstrated its enrichment at division septa, confirming its direct involvement in cell division. Deletion of *MAB_2363* resulted in attenuated virulence, as evidenced by reduced bacterial survival in macrophages and murine infection models. To assess its functional relation with *MAB_2362*, we compared the single-deletion mutant of *MAB_2363* with the single-deletion mutant of *MAB_2362* and the double-deletion mutant of *MAB_2362-MAB_2363*. Notably, the phenotypes of the *MAB_2363* mutant, including cell division defects, antibiotic susceptibility, and virulence, were markedly milder than those of the other two mutants. Collectively, these findings indicate that *MAB_2363* functions as a secondary but essential division-associated factor that operates during cell division, thereby influencing intrinsic resistance and virulence in *M. abscessus*.

## INTRODUCTION

*Mycobacterium abscessus* (*M. abscessus*), a rapidly growing nontuberculous mycobacterium, is a significant pathogen responsible for skin and soft tissue infections as well as pulmonary diseases primarily affecting immunocompromised individuals and patients with pre-existing pulmonary conditions (1, 2). Treatment of *M. abscessus* infections presents substantial clinical challenges owing to intrinsic resistance against multiple antibiotics, mediated by different mechanisms, including efflux pump systems, enzymatic modifications of drug targets or antimicrobial agents, and the highly impermeable cell wall (3). Current standard treatments involve prolonged (≥18 months) multidrug treatments (4–6), typically incorporating oral macrolides (e.g., azithromycin, clarithromycin) combined with parenteral aminoglycosides (e.g., amikacin) and β-lactam antibiotics (e.g., imipenem, cefoxitin), often supplemented by tetracyclines, oxazolidinones, or fluoroquinolones (7–9). Despite the use of these intensive multidrug regimens, cure rates ranged from 25% to 45% with elevated relapse frequencies (5, 6). These clinical limitations highlight the urgent need to deepen our understanding of *M. abscessus* drug-resistance mechanisms and to identify new therapeutic targets.

Cytokinesis in bacteria is coordinated by the divisome, a dynamic and multi-protein complex. The process is initiated by the GTP-dependent polymerization of the tubulin-like protein FtsZ into a Z-ring at the future division site, which subsequently recruits a cascade of proteins for septal peptidoglycan (PG) synthesis, hydrolysis, and remodeling (10, 11). A precise spatiotemporal balance between PG synthesis (by penicillin-binding proteins) and hydrolysis (by hydrolases such as peptidoglycan amidases and endopeptidases) is paramount for efficient daughter cell separation and the maintenance of cellular integrity (12, 13). Perturbation of this balance, as demonstrated in *Escherichia coli* and *Staphylococcus aureus*, results in profound morphological defects and often alters susceptibility to antibiotics (11, 14).

In Actinobacteria, including mycobacteria, cytokinesis is further complicated by their multi-layered cell envelope comprising a thick PG layer covalently linked to arabinogalactan and an outer membrane of mycolic acids (15, 16). Several factors that couple PG remodeling to envelope biogenesis have been identified in related species. For instance, depleting the divisome protein Wag31 (DivIVA) in *Mycobacterium smegmatis* (*M. smegmatis*) causes branched cells and division defects (17, 18), while loss of the PG amidase Ami1 in *Mycobacterium tuberculosis* (*M. tuberculosis*) results in cleavage defects and enhanced efficacy of β-lactams (19).

A conserved system exemplifying this coordination is the SteA-SteB system, first characterized in *Corynebacterium glutamicum* (*C. glutamicum*) (20). SteA and SteB localize to the division septum and are proposed to synchronize septal PG hydrolysis with the biogenesis of the outer layers. Deletion of either *steA* or *steB* leads to dramatic cytokinesis defects, along with a marked increase in susceptibility to the cell envelope-targeting antibiotic ethambutol (20). Homologs of this system exist in *M. abscessus*. Although it has been demonstrated that MAB_2362 (SteA) regulates cell division, thereby influencing intrinsic antibiotic resistance and virulence (21), the functional roles of MAB_2363, a candidate for SteB, in linking cell division to antibiotic resistance and virulence in this pathogen remain entirely unexplored.

In this study, we provide the first evidence that *MAB_2363* links cell division to intrinsic drug resistance and virulence in *M. abscessus*. Our findings reveal that MAB_2363 acts as a septum-localized division factor to maintain envelope integrity, antibiotic resistance, and pathogenicity. These discoveries advance our understanding of intrinsic resistance mechanisms and identify MAB_2363 as a promising target for therapeutic intervention.

## RESULTS

### Deletion of *MAB_2363* resulted in hypersensitivity of *M. abscessus* to multiple antibiotics *in vitro*

BLAST analysis showed that MAB_2363 shares 30.98% amino acid identity with SteB from *C. glutamicum* (Figure S1A). Despite the modest sequence identity, predicted 3D structure superimposition revealed a low root mean square deviation of 0.969 Å between MAB_2363 and SteB (Figure S2), indicating strong conformation similarity and supporting MAB_2363 as a SteB-like protein likely involved in cell division.

To investigate its functional role, we generated a *MAB_2363* knockout strain (Δ2363) using CRISPR-Cpf1-assisted non-homologous end joining. Sanger sequencing confirmed a 4-bp deletion causing a frameshift mutation (Figure S3A). Drug susceptibility testing revealed that Δ2363 was significantly more susceptible to multiple classes of antibiotics, including the protein synthesis inhibitors linezolid (LZD) and clarithromycin (CLA), the RNA polymerase inhibitor rifabutin (RFB), the β-lactam cefoxitin (CFX), the fluoroquinolones levofloxacin (LEV) and moxifloxacin (MXF), and the ATP synthase inhibitor bedaquiline (BDQ) (Table 1). Consistent results were obtained on solid medium (Figure 1 A). To determine the functional relationship between *MAB_2362* and *MAB_2363*, we compared Δ2363 with the previously characterized Δ2362 mutant and generated a Δ2362-2363 double knockout along with their complemented strains (Figure S3B). Across all tested antibiotics, Δ2363 consistently showed milder increases in susceptibility, whereas Δ2362 and Δ2362-2363 exhibited nearly identical minimum inhibitory concentrations (MICs), which were lower than those of Δ2363 (Table 1, Figure 1). These results indicate that *MAB_2362* plays a dominant role in modulating intrinsic drug resistance, whereas *MAB_2363* likely acts as a secondary factor. Drug susceptibility in all complemented strains was restored to near wild-type (WT) levels, confirming that the observed phenotypes were associated with the gene deletions. Growth curves of all mutants were comparable to WT (Figure S4), indicating that the increased antibiotic susceptibility is not due to growth defects.

**Figure 1.**
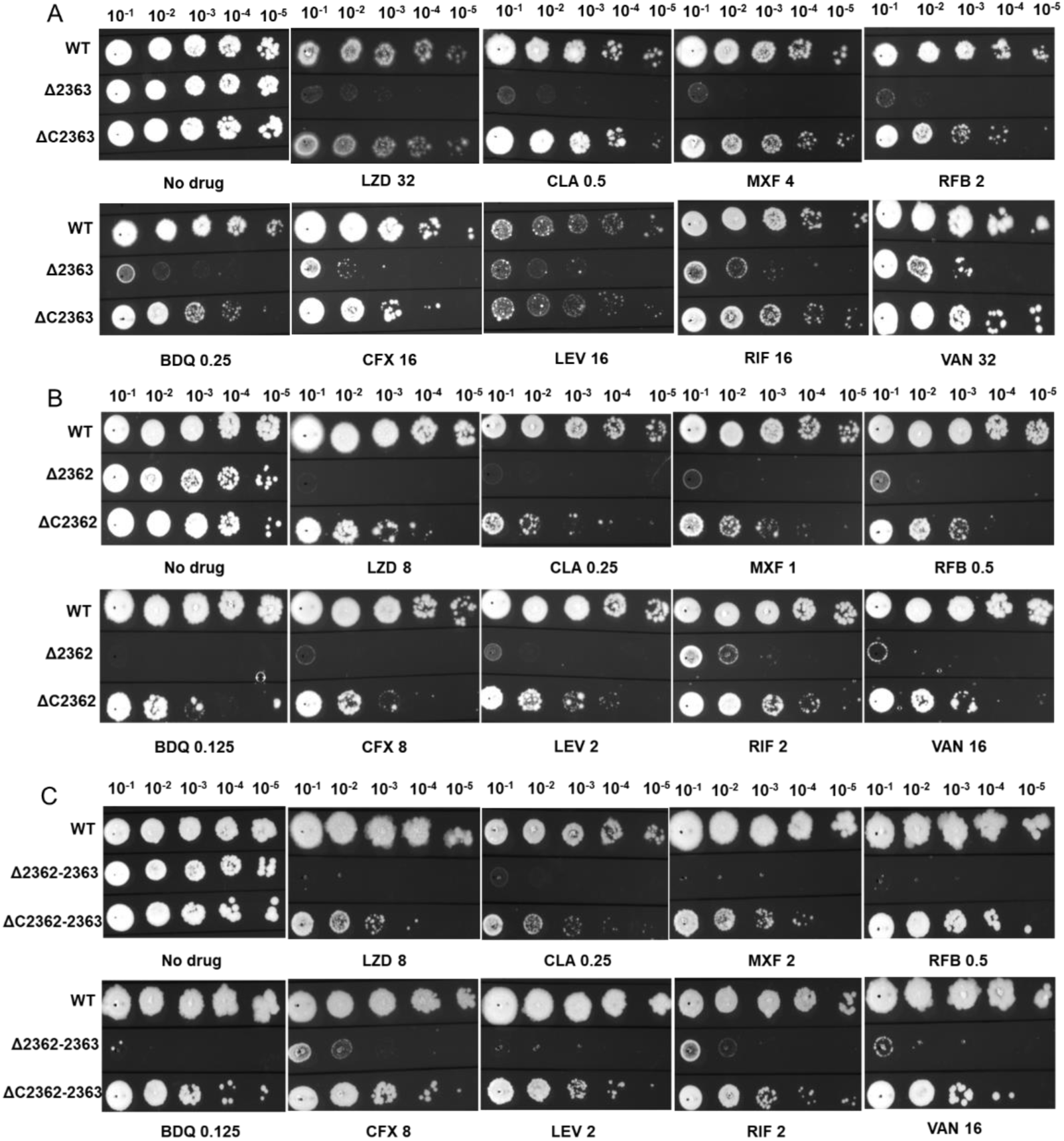
Susceptibility of various *M. abscessus* strains to indicated antibiotics on solid medium. WT, wild-type *M. abscessus*; Δ2363, *MAB_2363* knockout strain; ΔC2363, *MAB_2363* complemented strain; Δ2362, *MAB_2362* knockout strain; ΔC2362, *MAB_2362* complemented strain; Δ2362-2363, *MAB_2362*-*MAB_2363* double knockout strain; ΔC2362-2363, *MAB_2362*-*MAB_2363* double complemented strain.

**Table 1.**
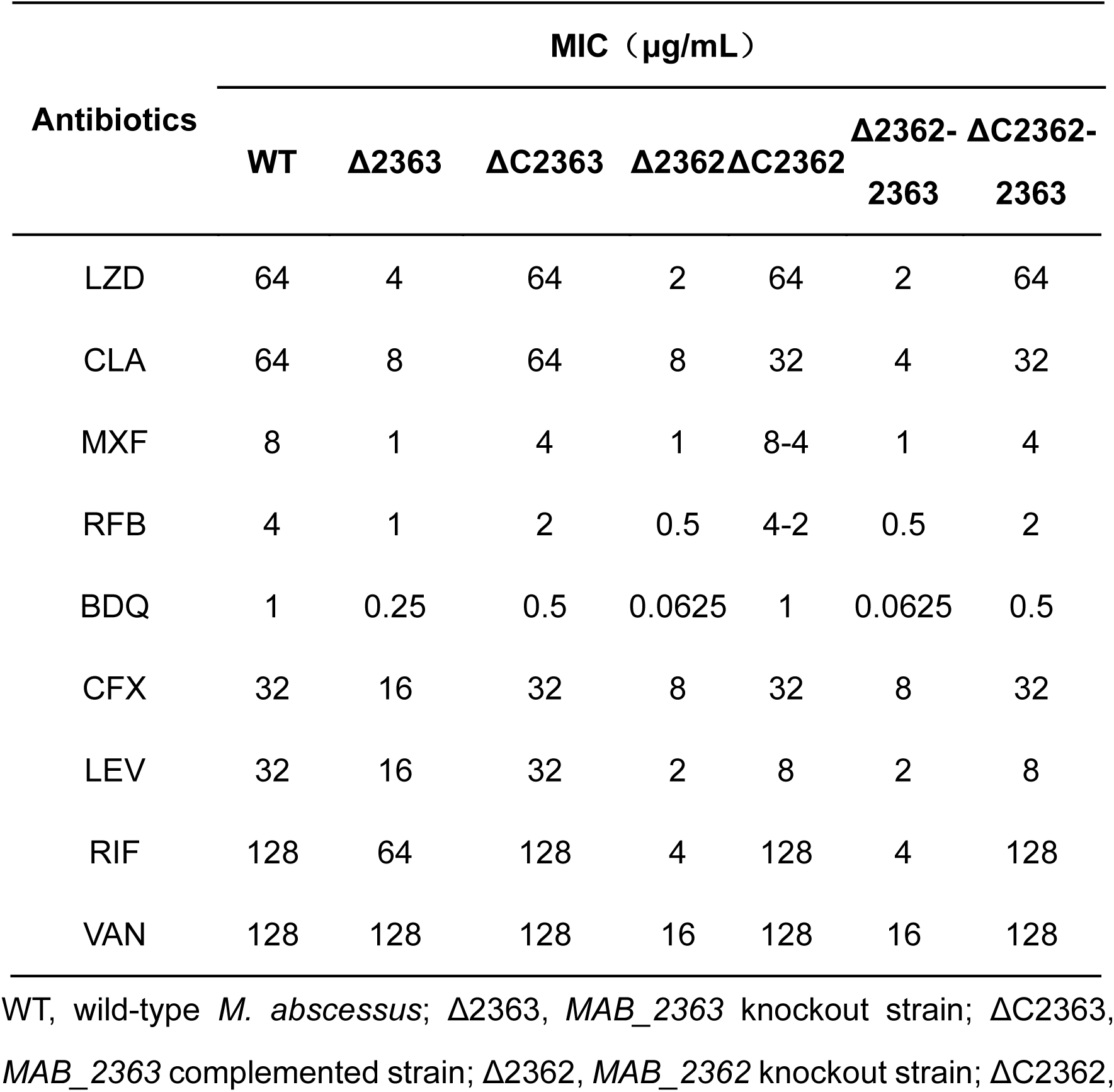
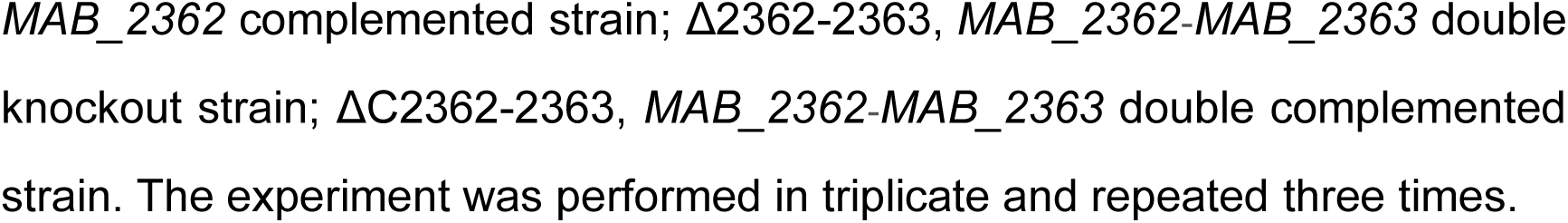
MICs of indicated antibiotics against various *M. abscessus* strains.

### Deletion of *MAB_2363* resulted in an increased cell envelope permeability of *M. abscessus*

The enhanced susceptibility of Δ2363 to multiple antibiotic classes suggested potential alterations in cell envelope integrity. To test this, we performed ethidium bromide (EtBr) accumulation assay.

The strain Δ2363 accumulated more EtBr than WT within 60 minutes (Figure 2), indicating increased cell wall permeability. However, the increase in Δ2363 was less pronounced than in Δ2362, whereas Δ2362 and Δ2362-2363 exhibited similar permeability levels, consistent with their respective antibiotic susceptibility profiles. These results indicate that *MAB_2363* contributes to maintaining cell envelope integrity in *M. abscessus*.

**Figure 2.**
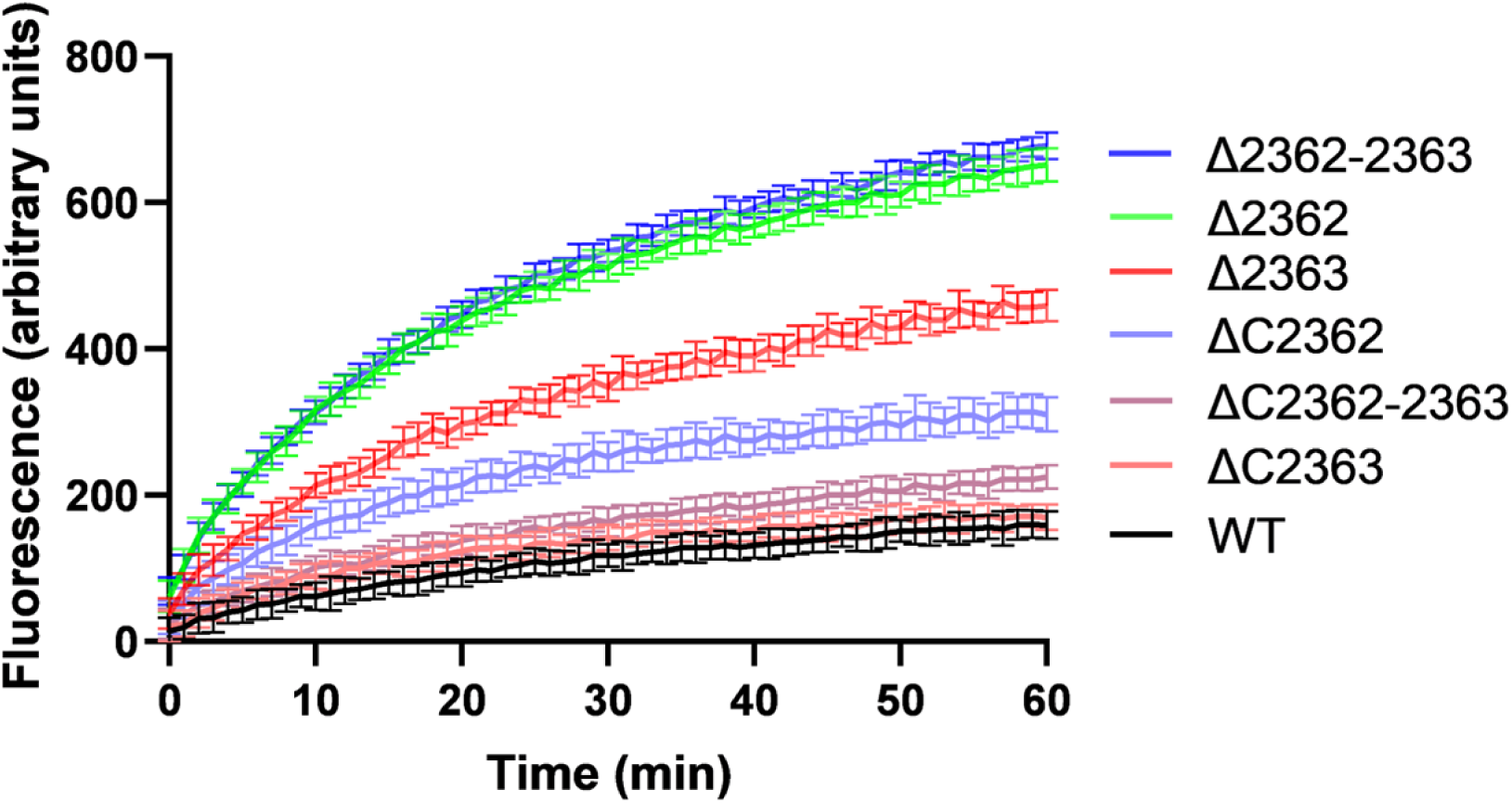
Cell wall permeability assay of various *M. abscessus* strains. WT, wild-type *M. abscessus*; Δ2363, *MAB_2363* knockout strain; ΔC2363, *MAB_2363* complemented strain; Δ2362, *MAB_2362* knockout strain; ΔC2362, *MAB_2362* complemented strain; Δ2362-2363, *MAB_2362*-*MAB_2363* double knockout strain; ΔC2362-2363, *MAB_2362*-*MAB_2363* double complemented strain. The experiment was repeated three times.

### Deletion of *MAB_2363* showed increased cell length and septa in *M. abscessus*

To investigate whether altered drug susceptibility and cell envelope integrity were associated with defects in cell division, we examined cellular morphology. Scanning electron microscopy revealed that Δ2363 cells were markedly elongated compared with WT, with a mean length of 2.45 µm versus 1.85 µm in WT (Figure 3 A-B, Table S1). Cells of Δ2362 and the Δ2362-2363 double mutant were even longer, measuring 3.24 µm and 3.66 µm, respectively. Complementation partially restored cell length in all strains (ΔC2363, ΔC2362, ΔC2362-2363: 2.17 µm, 2.08 µm, and 2.30 µm, respectively). Fluorescent D-amino acid HADA staining further revealed increased septation in Δ2363 (Figure 3 C-D). Cells with two or more septa accounted for 62% in Δ2363, compared with 11% in WT. Δ2362 and Δ2362-2363 exhibited similar septation frequencies (61% and 57%, respectively), while complementation largely restored septa numbers to near-WT levels. These results demonstrate that *MAB_2363* plays a significant role in regulating cell division, although its effects are milder than those of *MAB_2362*.

**Figure 3.**
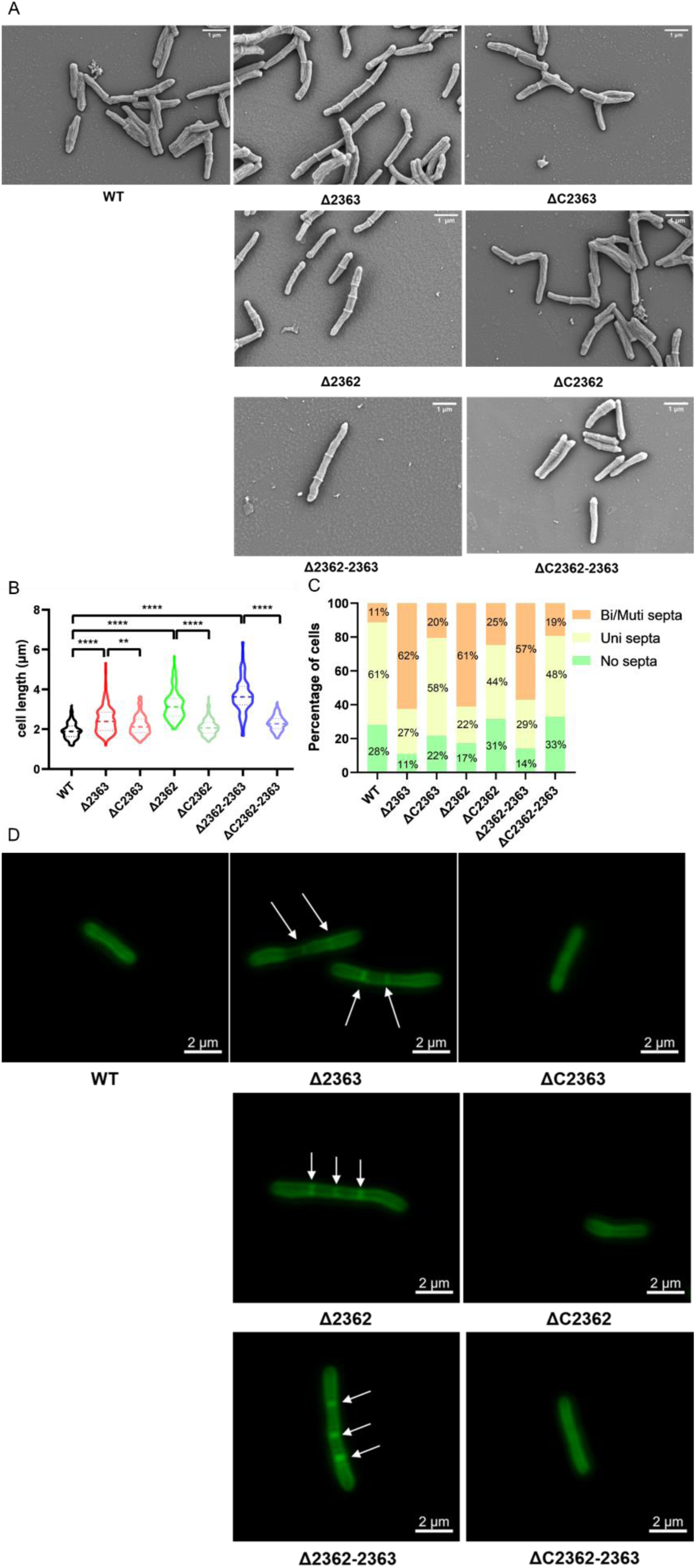
Microscopic analysis of cell length and septa in *M. abscessus* strains. (A) Representative scanning electron microscopy (SEM) images of different strains. (B) Violin plots comparing cell length among the seven strains. Statistical significance was determined by unpaired *t*-test; **, *P* < 0.01; ****, *P* < 0.0001. (C) Proportion of cells with different numbers of septa in each strain. (D) Representative laser confocal microscopy images of different *M. abscessus* strains. White arrows indicate cellular septa. WT, wild-type *M. abscessus*; Δ2363, *MAB_2363* knockout strain; ΔC2363, *MAB_2363* complemented strain; Δ2362, *MAB_2362* knockout strain; ΔC2362, *MAB_2362* complemented strain; Δ2362-2363, *MAB_2362*-*MAB_2363* double knockout strain; ΔC2362-2363, *MAB_2362*-*MAB_2363* double complemented strain.

### MAB_2363 was localized to the division septum in *M. abscessus*

To examine whether MAB_2363 directly participates in cell division, we assessed its subcellular localization using a GFP fusion. Fluorescence microscopy showed that GFP-MAB_2363 predominantly localized to the division septum, the site of new cell wall synthesis during cytokinesis (Figure 4). For comparison, GFP-MAB_2362 also localized to the septum. This septal localization mirrors that of established division regulators such as FtsZ and Wag31 (22, 23), supporting a direct role for MAB_2363 in cell division in *M. abscessus*.

**Figure 4.**
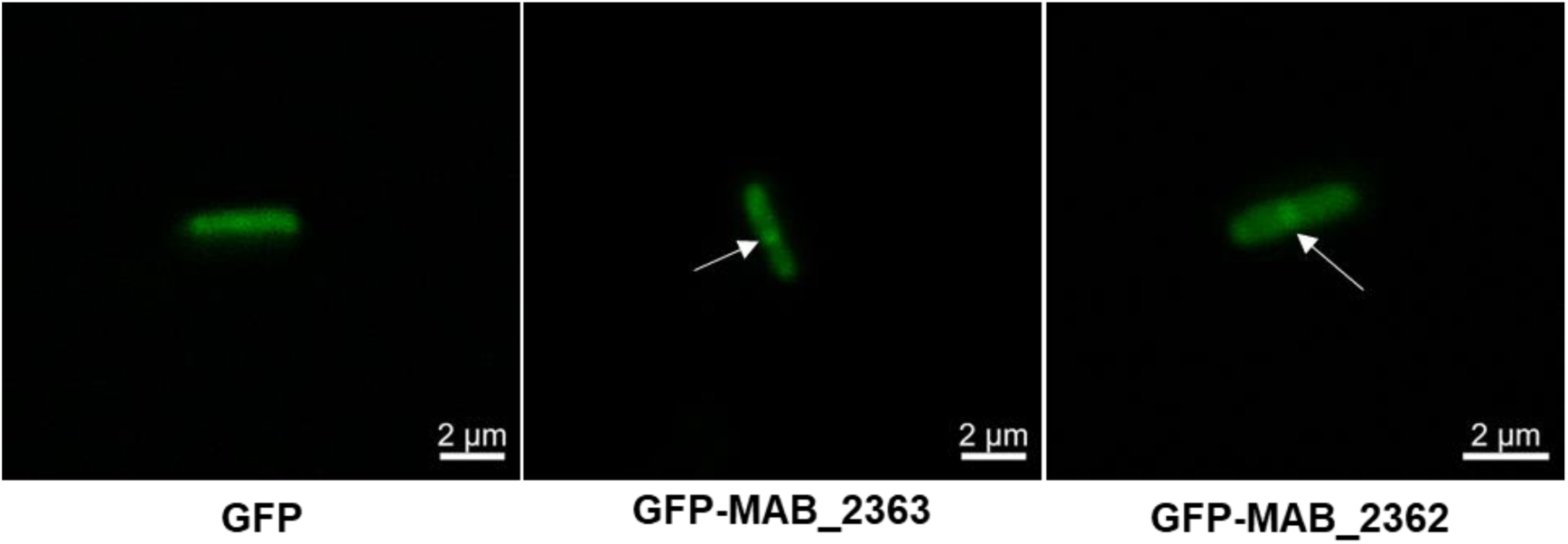
Subcellular localization of GFP fusion proteins in *M. abscessus*. Representative confocal fluorescence microscopy images of the indicated strains. White arrows mark septal localization.

### Deletion of *MAB_2363* increased *M. abscessus* susceptibility to BDQ *in vivo*

The MICs of BDQ and LZD against Δ2363 were 0.25 µg/mL and 4 µg/mL, respectively (Table 1), values predictive of potential *in vivo* efficacy (24–26). To test this, mice infected with WT or Δ2363 were treated with BDQ or LZD for 10 days. In WT-infected mice, neither drug reduced lung bacterial loads compared with untreated controls (Figure 5). In contrast, BDQ treatment significantly reduced lung colony forming units (CFUs) in Δ2363-infected mice, whereas LZD produced no measurable effect (Figure 5, Table S2). The lack of an observable response to LZD may reflect the attenuated virulence of the Δ2363 mutant, which results in a progressive, intrinsic decline in bacterial load over time and could obscure any additional impact of this bacteriostatic agent (27). These results indicate that deletion of *MAB_2363* enhances *M. abscessus* susceptibility to BDQ *in vivo*, hence presenting MAB_2363 as a critical regulator of intrinsic drug resistance and a potential target for novel therapeutics.

**Figure 5.**
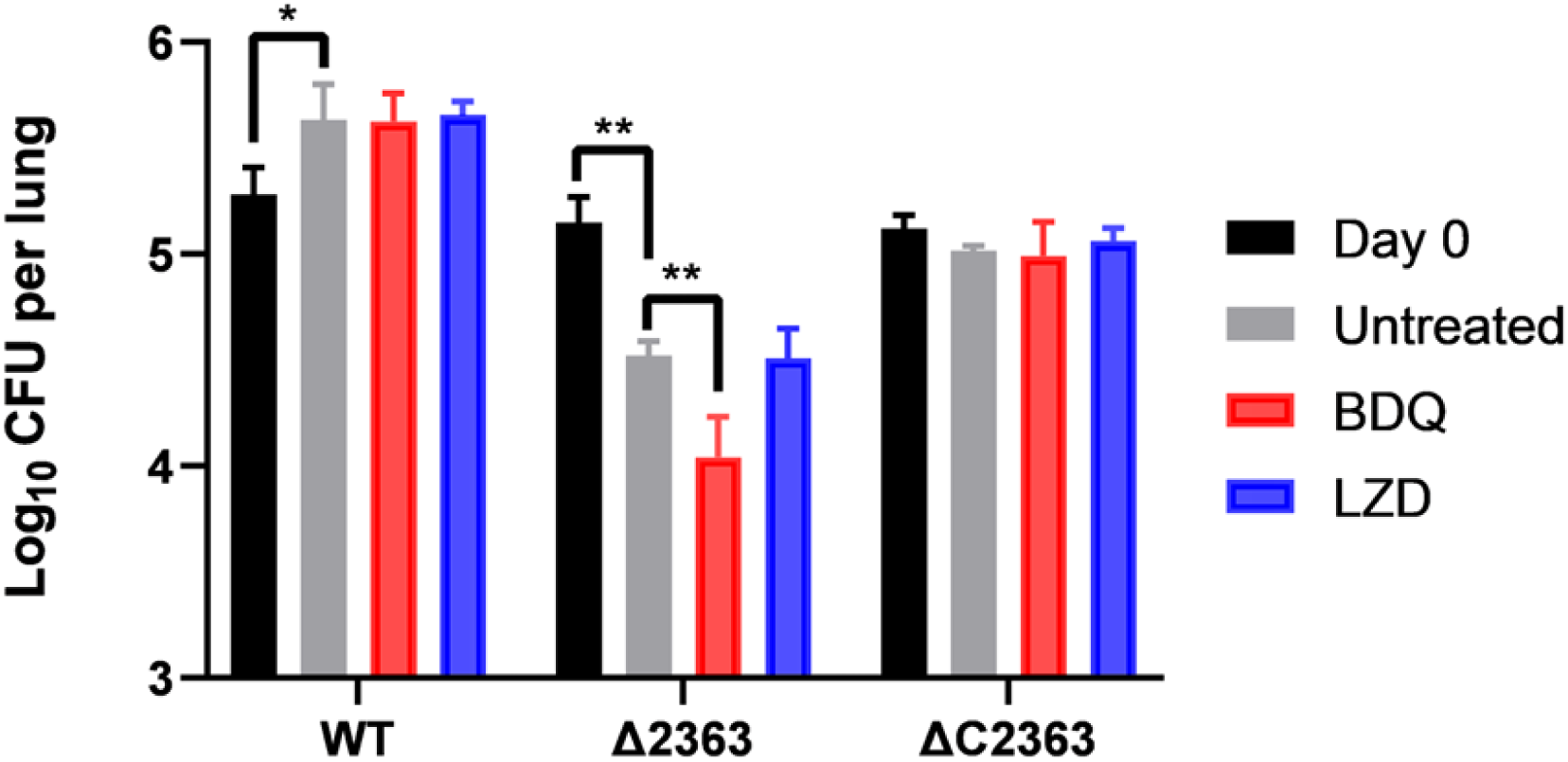
Assessment of susceptibility of different *M. abscessus* strains to BDQ and LZD *in vivo*. Lung bacterial burden over the 10-day treatment period. Drug doses (mg/kg): BDQ 20, LZD 100. Statistical significance was determined by unpaired *t*-test; *, *P* < 0.05; **, *P* < 0.01. WT, wild-type *M. abscessus*; Δ2363, *MAB_2363* knockout strain; ΔC2363, *MAB_2363* complemented strain.

### Deletion of *MAB_2363* significantly diminished the virulence of *M.* abscessus

Initial *in vivo* evaluation of BDQ and LZD revealed that, in untreated mice, pulmonary bacterial loads increased significantly in WT-infected mice by day 11 (*P* ≤ 0.05), whereas Δ2363-infected mice showed a decline (*P* ≤ 0.01), suggesting reduced virulence. To directly assess pathogenicity, we conducted follow-up experiments in macrophage and murine infection models. In RAW264.7 macrophages, Δ2363 exhibited significantly decreased intracellular survival at 72 hours post-infection compared with WT (Figure 6 A), indicating that deletion of *MAB_2363* impairs intramacrophage survival.

**Figure 6.**
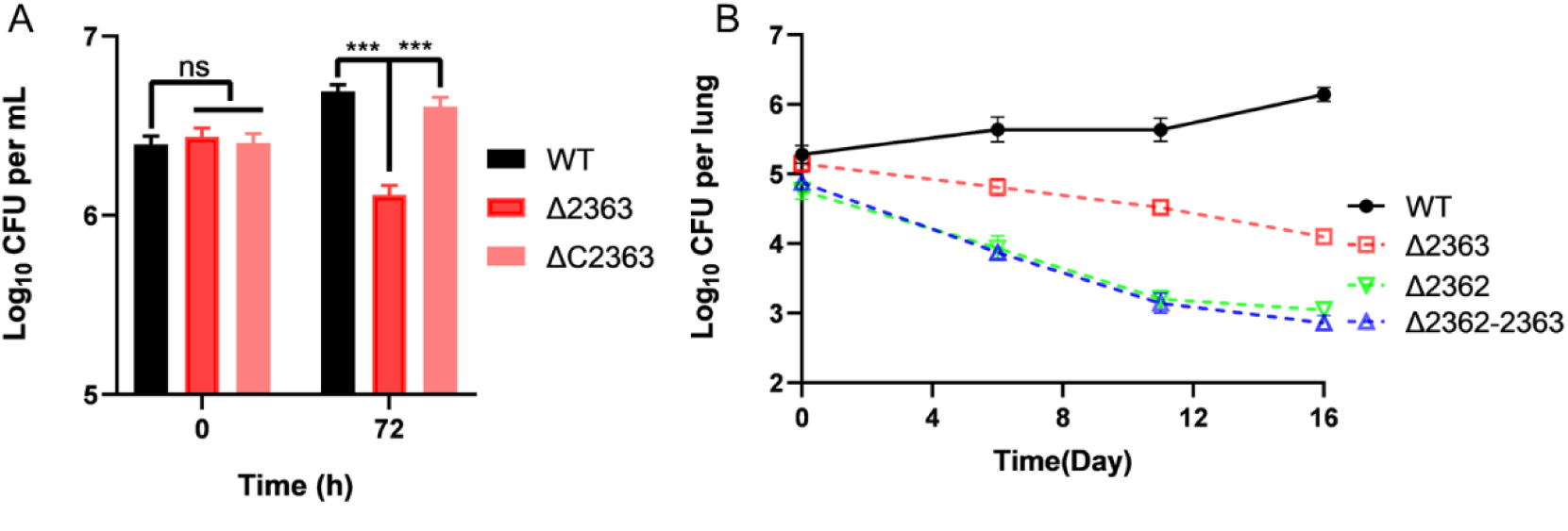
The virulence assessment of different *M. abscessus* strains. (A) Intracellular growth of different strains in macrophages. (B) Growth curves of different strains in a murine model in lungs. Statistical significance was determined by unpaired *t*-test; ns, nonsignificant; ***, *P* < 0.001. WT, wild-type *M. abscessus*; Δ2363, *MAB_2363* knockout strain; ΔC2363, *MAB_2363* complemented strain; Δ2362, *MAB_2362* knockout strain; Δ2362-2363, *MAB_2362*-*MAB_2363* double knockout strain.

To determine whether Δ2363 is attenuated *in viv*o, we monitored bacterial burdens at multiple time points over a 16-day murine infection course. WT bacterial loads increased steadily from 5.28 ± 0.13 to 6.14 ± 0.10 Log_10_ CFU per lung. In striking contrast, Δ2363 displayed a clear and consistent downward trajectory across all measured time points, ultimately declining from 5.15 ± 0.12 to 4.10 ± 0.03 Log_10_ CFU per lung by day 16 (Figure 6 B, Table S3). This sustained reduction, approximately 1 Log_10_, demonstrates that *MAB_2363* is required for maintaining bacterial burden during infection. Although Δ2362 and Δ2362-2363 exhibited even steeper declines (∼2 Log_10_), the consistent reduction observed for Δ2363 in macrophage and mouse models firmly establishes *MAB_2363* as an important determinant of *M. abscessus* virulence. The Δ2362 and Δ2362-2363 strains showed even greater reductions in lung bacterial burden (∼2 Log_10_), which is consistent with their more pronounced cell-division defects, that may further increase susceptibility to host clearance.

### Deletion of *MAB_2363* compromised tolerance to diverse environmental stresses

Given the reduced intracellular survival and attenuated virulence of Δ2363, we further evaluated the tolerance of Δ2363 to external stresses. Deletion of *MAB_2363* leads to impaired survival under surfactant, acidic, and oxidative conditions. Following a 4-hour incubation, Δ2363 exhibited significantly diminished survival rates compared to WT when exposed to SDS, acidic pH, or H_2_O_2_ (Figure 7). These results demonstrate that *MAB_2363* is required for *M. abscessus* to diverse environmental stresses likely encountered during infection and further underscore its role in maintaining envelope integrity and stress resilience.

**FIG 7.**
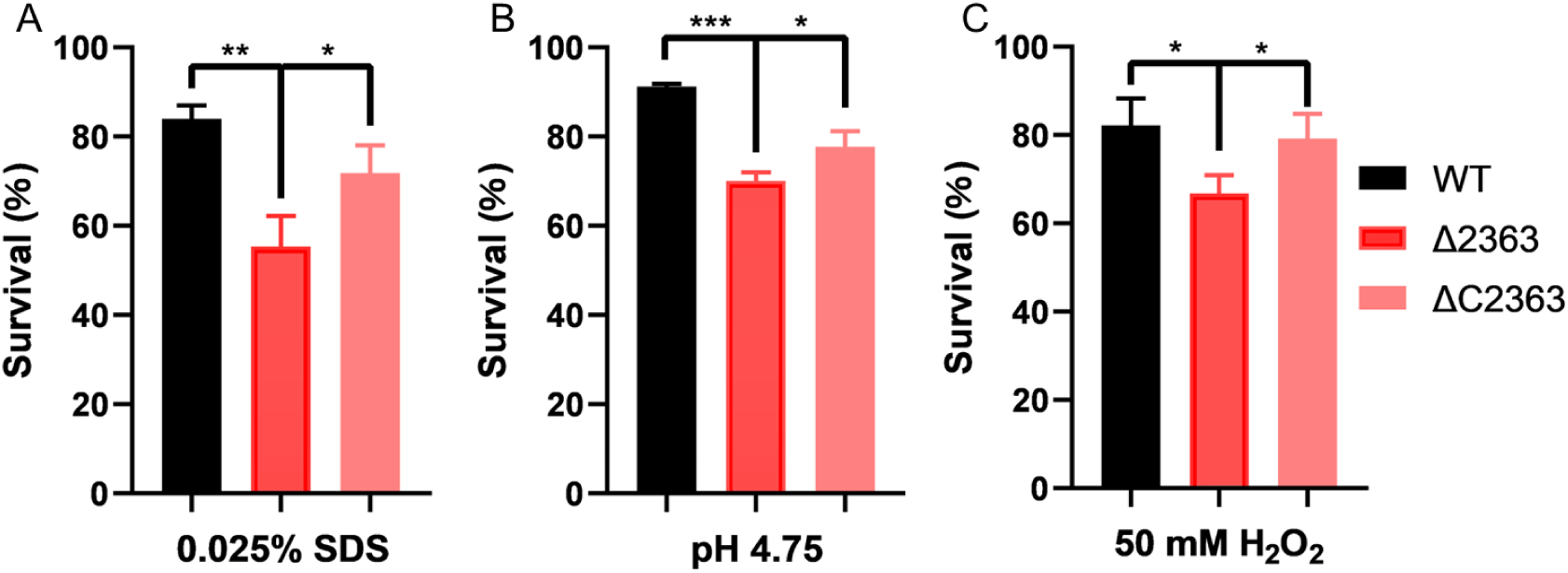
Survival of different *M. abscessus* strains under diverse stresses. (A) SDS stress: Strains were cultured in medium supplemented with 0.025% SDS. (B) Acid stress: Strains were exposed to medium adjusted to pH 4.75. (C) Oxidative stress: Strains were grown in medium containing 50 mM H_2_O_2_. Survival rates were evaluated by measuring CFU/mL at 4 hours. Statistical significance was determined by unpaired *t*-test; ns, nonsignificant; *, *P* < 0.05; **, *P* < 0.01; ***, *P* < 0.001. WT, wild-type *M. abscessus*; Δ2363, *MAB_2363* knockout strain; ΔC2363, *MAB_2363* complemented strain. The experiment was repeated three times.

## DISCUSSION

*M. abscessus,* a notoriously drug-resistant nontuberculous mycobacterial pathogen, causes severe pulmonary diseases and skin or soft tissue infections, particularly in immunocompromised individuals (1, 2). Management of these infections remains clinically challenging due to their intrinsic resistance to most frontline antibiotics (3). Identifying genetic determinants that underlie this intrinsic resistance is therefore essential for developing new therapeutic strategies or potentiating the activity of existing drugs. In this study, we demonstrate that deletion of *MAB_2363* markedly sensitizes *M. abscessus* to multiple antibiotic classes, revealing a previously unrecognized determinant of intrinsic multidrug resistance.

The broad-spectrum sensitization of Δ2363 across multiple antibiotics suggests that *MAB_2363* plays a role in maintaining overarching cellular integrity rather than modulating distinct drug targets. Consistent with this interpretation, Δ2363 exhibited increased cell-envelope permeability and pronounced morphological abnormalities. These phenotypes align with those reported for other envelope- or division-associated proteins in actinomycetes. For example, depletion of Wag31 in *M. smegmatis* reduces resistance to lipophilic antibiotics by compromising envelope integrity (28), and loss of CwsA disrupts PG synthesis and cell division (29). Similarly, disruption of the SteA-SteB system in *C. glutamicum* results in elongated cells, abnormal septum formation, and enhanced susceptibility to cell wall-active antibiotics (20). Although MAB_2363 shares limited amino acid sequence identity with SteB from *C. glutamicum*, their high structural similarity supports a conserved functional role in division-associated envelope maintenance. Thus, the increased permeability observed in Δ2363 provides a consistent mechanistic explanation for its heightened susceptibility to diverse antibiotics.

Our findings further suggest that *MAB_2363* is required for proper cell division. Δ2363 cells were elongated and exhibited increased septation, and GFP-MAB_2363 localized to the division septum. In mycobacteria, septal remodeling and envelope assembly are tightly coordinated processes, therefore, defects in division frequently translate into envelope instability (30, 31). The milder defects observed in Δ2363 relative to Δ2362 correspond to their differential impacts on permeability and antibiotic susceptibility, suggesting that the two proteins play related yet distinct roles during cytokinesis.

We additionally demonstrate that loss of MAB_2363 attenuates *M. abscessus* virulence in macrophage and murine infection models. The declining bacterial burden observed in Δ2363-infected mice likely reflects its compromised envelope integrity, rendering Δ2363 less capable of resisting host-derived stresses such as oxidative damage and acidic conditions. Similar virulence attenuation has been reported for mutants with envelope defects, including those lacking LmeA or inhibited in Ag85C activity (32, 33). The attenuation of Δ2363 is therefore consistent with its structural defects and further highlights the role of envelope stability in pathogenic fitness.

Functionally, MAB_2363 is annotated as a copper transporter (NCBI GenBank accession number WP_005058615.1), a protein implicated in copper ion transport and homeostasis. Although copper is essential for numerous enzymatic processes, excessive accumulation is toxic. In *M. tuberculosis* and *M. smegmatis*, the copper transporter MctB homologs encoded by *Rv1698* and *MSMEG_3747* help maintain copper homeostasis, and their deletion confers copper hypersensitivity (34). However, loss of MAB_2363 in *M. abscessus* did not exhibit copper sensitivity (Figure S5). Moreover, heterologous expression of *Rv1698* or *MSMEG_3747* in the Δ2363 background failed to restore antibiotic resistance (Table S4). Despite moderate amino acid conservation with these homologs (55.77% and 58.01%, respectively; Figure S1B), MAB_2363 did not functionally substitute for them, indicating divergence in biological roles. Supporting this, GFP fusion imaging showed that MAB_2363 predominantly localizes to the division septum rather than displaying a distribution typical of copper transporters (Figure 4). Collectively, these findings indicate that, despite sequence conservation, MAB_2363 has acquired a species-specific function linked to cell division-associated processes rather than copper homeostasis.

In summary, this study reveals MAB_2363 as an important determinant of envelope integrity, cell division, intrinsic antibiotic resistance, and virulence in *M. abscessus*. Disruption of MAB_2363 markedly enhances susceptibility to multiple antibiotics, including the clinically important agent BDQ, and reduces bacterial survival *in vivo*. Taken together, these findings highlight *MAB_2363* as a promising therapeutic target: inhibiting its function may weaken the cell envelope, potentiate the efficacy of existing drugs, and attenuate pathogenicity, collectively providing a compelling strategy for improving treatment outcomes against *M. abscessus* infections.

## MATERIALS AND METHODS

### Strains, plasmids and culture conditions

*M. abscessus* GZ002 (NCBI GenBank accession number CP034181), was obtained from Guangzhou Chest Hospital (35, 36). The derivative strain Mab: pNHEJ-Cpf1 was constructed previously in our laboratory (37). Mycobacteria were cultured in Middlebrook 7H9 broth (Difco) medium supplemented with 0.2% glycerol, 0.05% Tween-80, and 10% OADC or on Middlebrook 7H10/7H11 agar supplemented with 0.5% glycerol and 10% OADC. Medium was further supplemented with kanamycin (KAN, 100 µg/mL) or zeocin (ZEO, 30 µg/mL) as required. Anhydrotetracycline (ATc) was used at 200 ng/mL for induction. *Escherichia coli* DH5α was grown in LB broth or on LB agar supplemented with KAN (50 µg/mL) or ZEO (30 µg/mL) at 37°C. The strains used in this study are listed in Table S5. RAW264.7 macrophages were cultured in Dulbecco’s Modified Eagle Medium (DMEM, Sbjbio) supplemented with 10% fetal bovine serum (FBS, Sbjbio) at 37°C under 5% CO_2_. Restriction enzymes *Bpm* I (New England Biolabs), *Hin*d III, *Cla* I, and *Bam*H I (TaKaRa) were commercially sourced. Plasmids used in this study are listed in Table S6.

### Construction of knockout and complemented strains

The genes were knocked out using a CRISPR-Cpf1-assisted non-homologous end joining system (37). Guide RNAs were designed, synthesized, annealed, and cloned into linearized pCR-Zeo plasmids. The recombinant plasmids were electroporated into the Mab:pNHEJ-Cpf1. Transformants were selected on 7H11 agar plates supplemented with KAN (100 µg/mL), ATc (200 ng/mL), and ZEO (30 µg/mL) at 30°C for 5 days. The knockout strains were identified by screening single colonies using PCR, and confirmed by Sanger sequencing.

For complementation, the target gene was amplified and cloned into the expression vector pMV261 via Gibson assembly. The resulting construct was then introduced into the knockout strain by electroporation to generate the complemented strain. The GFP fusion strain was constructed using the same strategy. All plasmids and PCR primers used in this study are listed in Table S6 and Table S7.

### Antibiotic susceptibility testing

The minimum inhibitory concentrations (MICs) of different antibiotics were determined using a broth microdilution method. Bacterial cultures were grown to an optical density (OD_600_) of 0.6-0.8. Twofold serial dilutions of antibiotics were prepared in a 96-well plate, and bacteria were diluted to a final concentration of 5 × 10^5^ CFUs/mL in Tween-80-free 7H9 medium. The plates were incubated at 37°C for 3 days except for CLA, which required a 14-day incubation (38). Experiments were performed in triplicate. The MIC was defined as the lowest antibiotic concentration that completely inhibited visible bacterial growth. To assess antibiotic susceptibility on solid media, strains were cultured to the exponential growth phase. A 10-fold serial dilution of bacterial suspensions was prepared, and 2 μL aliquots were spotted onto 7H10 agar plates containing different antibiotic concentrations or no antibiotic. Plates were incubated at 37°C for 3 days.

### Cell wall permeability assay

As described previously, the cell wall permeability of strains was evaluated using EtBr uptake assays (25). Bacterial cultures were grown to mid-log phase, washed three times with PBS containing 0.05% Tween 80, and resuspended in the same buffer to an OD_600_ of 0.5. An EtBr solution (4 μg/mL) was prepared in PBS with 0.05% Tween 80, supplemented with glucose to a final concentration of 0.8%. Aliquots (100 μL) of this solution were transferred to a white 96-well plate, followed by the addition of 100 μL of bacterial suspension. Fluorescence of EtBr was measured every minute for 60 minutes using a FlexStation 3 (Molecular Devices, USA) with excitation and emission wavelengths of 530 nm and 590 nm, respectively. All experiments were conducted in triplicate.

### Microscopic analysis

Strains were cultured to an OD_600_ of 0.6-0.8. Subsequently, 1 mL of the bacterial suspension was centrifuged and fixed with 4% glutaraldehyde at room temperature for 1 h. After centrifugation, the supernatant was discarded, and the cells were washed three times with PBS before resuspension. 10 μL aliquot of the bacterial suspension was transferred to a circular glass substrate and air-dried at room temperature, followed by three times PBS washes. Samples were then dehydrated through washing with graded series of ethanol concentrations (70%, 80%, 90%, and 100%). Critical point drying was conducted to preserve their innate morphology and obtain high-quality images. Afterward, samples were sputter-coated with gold prior to imaging using a field-emission scanning electron microscope (GeminiSEM, Zeiss). Cell lengths were determined using Nano Measurer 1.2.

For peptidoglycan labeling, strains were grown to an OD_600_ of 0.6-0.8 and diluted to an OD_600_ of 0.3 using 7H9 medium. Cells were then incubated in medium containing 1 mM HADA (7-hydroxy-9H-(1,3-dichloro-9,9-dimethylacridin-2-one)) for 4 h under light-protected conditions. After incubation, bacterial pellets were washed three times with PBS containing 0.05% Tween 80 and fixed with 4% paraformaldehyde for 30 mins. Fluorescently labeled samples were visualized using a laser scanning confocal microscope (LSM710, Zeiss).

To localize GFP-tagged proteins, *gfp* and *MAB_2363* gene fragments were cloned into a linearized pMV261 plasmid via three-fragment recombination to generate the plasmid pMV-GFP-MAB_2363. Similarly, *gfp* was recombined with the linearized pMV261 plasmid to construct pMV-GFP. These plasmids were electroporated into WT, yielding Mab:GFP-MAB_2363 and Mab:GFP. The Mab:GFP-MAB_2363 strain expressed the GFP-MAB_2363 fusion protein. Cultures were grown to an OD_600_ of 0.6–0.8, washed three times with PBS–0.05% Tween 80, fixed with 4% paraformaldehyde for 30 minutes, and visualized using a Zeiss LSM 710 Confocal Laser Scanning Microscope to observe MAB_2363 localization. The localization of MAB_2362 was analyzed following the same procedure.

### Assessment of intracellular bacterial survival

Bacterial strains were cultured to an OD_600_ of 0.6-0.8, washed three times with PBS, and diluted to a multiplicity of infection (MOI) of 10:1. The final bacterial suspension was adjusted to 1 × 10^5^ CFU/mL using DMEM. A 100 μL aliquot of the diluted suspension was added to a 96-well plate containing 100 μL adherent cells. After 4 h of incubation at 37°C with 5% CO_2_, cells were washed three times with DMEM to remove non-internalized bacteria. To eliminate extracellular bacteria, 200 μL of DMEM supplemented with 250 μg/mL amikacin was added, followed by 2 h incubation. Cells were then washed three times with DMEM and maintained in 200 μL of DMEM containing 50 μg/mL amikacin to suppress residual extracellular bacterial growth. Intracellular bacterial loads were quantified at 0 and 72 hours post-infection by lysing infected cells in sterile water, serially diluting lysates, and plating on 7H10 agar. Experiments were performed in triplicate.

### Animal experiments

Ethical approval was obtained from the institutional animal care and use committee of Guangzhou Institutes of Biomedicine and Health, Chinese Academy of Sciences (IACUC number 2025089). Female BALB/c mice, aged 6-8 weeks and weighing 19-22 grams (GuangDong GemPharmatech Co., Ltd.) were used in this study. To achieve effective immunosuppression, all mice received daily dexamethasone (DEXA; D1756, Sigma-Aldrich) treatment starting before infection and continuing throughout the experimental period. As previously described (39), DEXA was dissolved in sterile PBS and administered via subcutaneous injection at a dose of 4 mg/kg/day. Mice were infected with logarithmic phase cells of each of the bacterial strains via aerosol exposure. Five hours post-infection, five mice from each group were euthanized to determine the initial bacterial load in the lungs. Lungs were homogenized in PBS, serially diluted ten fold, and plated on 7H10 agar plates for CFU enumeration. On day 6, 11, and 16 post-infection, five mice per group were euthanized, and lungs were aseptically collected for CFU counting. In the drug-treated groups, mice were administered LZD (100 mg/kg) and BDQ (20 mg/kg) daily via oral gavage, starting on day 1 post-infection, for a total of 10 days. All treated mice were sacrificed on day 11. The lungs were collected, homogenized, and plated for bacterial enumeration. To eliminate drug carryover effects, homogenates from the BDQ-treated group were plated on 7H10 agar supplemented with 0.4% activated charcoal.

### Determination of bacterial survival under different environmental stresses

Bacterial strains were cultured to an OD_600_ of 0.6-0.8 and then washed three times with 7H9 medium. The OD_600_ was adjusted to 0.5, and the bacteria were cultured in media containing 0.025% SDS, 50 mM H_2_O_2_, or pH 4.75. Bacterial CFU was determined at 0 and 4 hours. All experiments were repeated three times.

## Supporting information

Clean Manuscript

## ACKNOWLEDGEMENTS

This work was supported by the National Key R&D Program of China (2021YFA1300904), National Natural Science Foundation of China (82502762, 82304575), Guangdong Provincial Basic and Applied Basic Research Fund (2024A1515012412), Guangzhou Science and Technology Plan-Youth Doctoral ‘Sail’ Project (2024A04J4273), the State Key Lab of Respiratory Disease, Guangzhou Institute of Respiratory Diseases, First Affiliated Hospital of Guangzhou Medical University (SKLRD-Z-202412, SKLRD-Z-202301, and SKLRD-OP-202324), Key Research and Development Program of Guangzhou (2025B01J3019), Guangzhou National Laboratory and State Key Laboratory of Respiratory Disease, GZNL2025B01006, and Major Project of Guangzhou National Laboratory, GZNL2025C01003).

## Notes

### Competing Interest Statement

The authors have declared no competing interest.

## References

1. M. R. Lee, W. H. Sheng, C. C. Hung, C. J. Yu, L. N. Lee and P. R. Hsueh. 2015. *Mycobacterium abscessus* complex infections in humans. Emerg Infect Dis 21:1638–1646. 10.3201/2109.141634

2. D. S. Prince, D. D. Peterson, R. M. Steiner, J. E. Gottlieb, R. Scott, H. L. Israel, W. G. Figueroa and J. E. Fish. 1989. Infection with *Mycobacterium avium* complex in patients without predisposing conditions. N Engl J Med 321:863–868. 10.1056/nejm198909283211304

3. S. Luthra, A. Rominski and P. Sander. 2018. The role of antibiotic-target-modifying and antibiotic-modifying enzymes in *Mycobacterium abscessus* drug resistance. Front Microbiol 9:2179. 10.3389/fmicb.2018.02179

4. D. E. Griffith, W. M. Girard and R. J. Wallace, Jr. 1993. Clinical features of pulmonary disease caused by rapidly growing mycobacteria. An analysis of 154 patients. Am Rev Respir Dis 147:1271–1278. 10.1164/ajrccm/147.5.1271

5. J. Jarand, A. Levin, L. Zhang, G. Huitt, J. D. Mitchell and C. L. Daley. 2011. Clinical and microbiologic outcomes in patients receiving treatment for *Mycobacterium abscessus* pulmonary disease. Clin Infect Dis 52:565–571. 10.1093/cid/ciq237

6. K. Jeon, O. J. Kwon, N. Y. Lee, B. J. Kim, Y. H. Kook, S. H. Lee, Y. K. Park, C. K. Kim and W. J. Koh. 2009. Antibiotic treatment of *Mycobacterium abscessus* lung disease: a retrospective analysis of 65 patients. Am J Respir Crit Care Med 180:896–902. 10.1164/rccm.200905-0704OC

7. R. A. Floto, K. N. Olivier, L. Saiman, C. L. Daley, J. L. Herrmann, J. A. Nick, P. G. Noone, D. Bilton, P. Corris, R. L. Gibson, S. E. Hempstead, K. Koetz, K. A. Sabadosa, I. Sermet-Gaudelus, A. R. Smyth, J. van Ingen, R. J. Wallace, K. L. Winthrop, B. C. Marshall and C. S. Haworth. 2016. US Cystic Fibrosis Foundation and European Cystic Fibrosis Society consensus recommendations for the management of non-tuberculous mycobacteria in individuals with cystic fibrosis: executive summary. Thorax 71:88–90. 10.1136/thoraxjnl-2015-207983

8. D. E. Griffith, T. Aksamit, B. A. Brown-Elliott, A. Catanzaro, C. Daley, F. Gordin, S. M. Holland, R. Horsburgh, G. Huitt, M. F. Iademarco, M. Iseman, K. Olivier, S. Ruoss, C. F. von Reyn, R. J. Wallace, Jr. and K. Winthrop. 2007. An official ATS/IDSA statement: diagnosis, treatment, and prevention of nontuberculous mycobacterial diseases. Am J Respir Crit Care Med 175:367–416. 10.1164/rccm.200604-571ST

9. C. S. Haworth, J. Banks, T. Capstick, A. J. Fisher, T. Gorsuch, I. F. Laurenson, A. Leitch, M. R. Loebinger, H. J. Milburn, M. Nightingale, P. Ormerod, D. Shingadia, D. Smith, N. Whitehead, R. Wilson and R. A. Floto. 2017. British Thoracic Society guidelines for the management of non-tuberculous mycobacterial pulmonary disease (NTM-PD). Thorax 72:ii1–ii64. 10.1136/thoraxjnl-2017-210927

10. S. S. Shell, M. Bar-Oz, J. Xiao, M. Pandey, J. Bellardinelli, O. I. Ibitoye, M. Jackson, S. H. Oehlers, D. Barkan and M. Meir. 2025. GplR1, an unusual TetR-like transcription factor in *Mycobacterium abscessus*, controls the production of cell wall glycopeptidolipids, colony morphology, and virulence. mSystems 10:e0087225. 10.1128/msystems.00872-25

11. A. Typas, M. Banzhaf, C. A. Gross and W. Vollmer. 2011. From the regulation of peptidoglycan synthesis to bacterial growth and morphology. Nat Rev Microbiol 10:123–136. 10.1038/nrmicro2677

12. S. Bhamidi, L. Shi, D. Chatterjee, J. T. Belisle, D. C. Crick and M. R. McNeil. 2012. A bioanalytical method to determine the cell wall composition of *Mycobacterium tuberculosis* grown *in vivo*. Anal Biochem 421:240–249. 10.1016/j.ab.2011.10.046

13. P. Seiler, T. Ulrichs, S. Bandermann, L. Pradl, S. Jörg, V. Krenn, L. Morawietz, S. H. Kaufmann and P. Aichele. 2003. Cell-wall alterations as an attribute of *Mycobacterium tuberculosis* in latent infection. J Infect Dis 188:1326–1331. 10.1086/378563

14. B. B. Aldridge, M. Fernandez-Suarez, D. Heller, V. Ambravaneswaran, D. Irimia, M. Toner and S. M. Fortune. 2012. Asymmetry and aging of mycobacterial cells lead to variable growth and antibiotic susceptibility. Science 335:100–104. 10.1126/science.1216166

15. C. Baranowski, E. H. Rego and E. J. Rubin. 2019. The dream of a mycobacterium. Microbiol Spectr 7(2):10.1128. 10.1128/microbiolspec.GPP3-0008-2018

16. K. J. Kieser and E. J. Rubin. 2014. How sisters grow apart: mycobacterial growth and division. Nat Rev Microbiol 12:550–562. 10.1038/nrmicro3299

17. N. H. Arejan, D. R. Czapski, J. A. Buonomo and C. C. Boutte. 2024. MmpL3, Wag31, and PlrA are involved in coordinating polar growth with peptidoglycan metabolism and nutrient availability. J Bacteriol 206:e0020424. 10.1128/jb.00204-24

18. C. M. Kang, S. Nyayapathy, J. Y. Lee, J. W. Suh and R. N. Husson. 2008. Wag31, a homologue of the cell division protein DivIVA, regulates growth, morphology and polar cell wall synthesis in mycobacteria. Microbiology (Reading) 154:725–735. 10.1099/mic.0.2007/014076-0

19. S. Senzani, D. Li, A. Bhaskar, C. Ealand, J. Chang, B. Rimal, C. Liu, S. Joon Kim, N. Dhar and B. Kana. 2017. An Amidase_3 domain-containing N-acetylmuramyl-L-alanine amidase is required for mycobacterial cell division. Sci Rep 7:1140. 10.1038/s41598-017-01184-7

20. H. C. Lim, J. W. Sher, F. P. Rodriguez-Rivera, C. Fumeaux, C. R. Bertozzi and T. G. Bernhardt. 2019. Identification of new components of the RipC-FtsEX cell separation pathway of *Corynebacterineae*. PLoS Genet 15:e1008284. 10.1371/journal.pgen.1008284

21. Y. Ju, L. Li, J. Zhang, B. Yusuf, S. Zeng, C. Fang, X. Tian, X. Han, J. Ding, H. Zhang, W. Ma, S. Wang, X. Chen and T. Zhang. 2025. The gene *MAB_*2362 is responsible for intrinsic resistance to various drugs and virulence in *Mycobacterium abscessus* by regulating cell division. Antimicrob Agents Chemother 69:e0043324. 10.1128/aac.00433-24

22. S. Du, S. Pichoff, K. Kruse and J. Lutkenhaus. 2018. FtsZ filaments have the opposite kinetic polarity of microtubules. Proc Natl Acad Sci U S A 115:10768–10773. 10.1073/pnas.1811919115

23. E. H. Rego, R. E. Audette and E. J. Rubin. 2017. Deletion of a mycobacterial divisome factor collapses single-cell phenotypic heterogeneity. Nature 546:153–157. 10.1038/nature22361

24. C. L. Daley, J. M. Iaccarino, C. Lange, E. Cambau, R. J. Wallace, Jr., C. Andrejak, E. C. Böttger, J. Brozek, D. E. Griffith, L. Guglielmetti, G. A. Huitt, S. L. Knight, P. Leitman, T. K. Marras, K. N. Olivier, M. Santin, J. E. Stout, E. Tortoli, J. van Ingen, D. Wagner and K. L. Winthrop. 2020. Treatment of nontuberculous mycobacterial pulmonary disease: an official ATS/ERS/ESCMID/IDSA clinical practice guideline. Eur Respir J 56(1):2000535. 10.1183/13993003.00535-2020

25. S. Wang, X. Cai, W. Yu, S. Zeng, J. Zhang, L. Guo, Y. Gao, Z. Lu, H. M. A. Hameed, C. Fang, X. Tian, B. Yusuf, C. Chhotaray, M. D. S. Alam, B. Zhang, H. Ge, D. A. Maslov, G. M. Cook, J. Peng, Y. Lin, N. Zhong, G. Zhang and T. Zhang. 2022. Arabinosyltransferase C mediates multiple drugs intrinsic resistance by altering cell envelope permeability in *Mycobacterium abscessus*. Microbiol Spectr 10:e0276321. 10.1128/spectrum.02763-21

26. J. Zhang, Y. Ju, L. Li, H. M. A. Hameed, B. Yusuf, Y. Gao, C. Fang, X. Tian, J. Ding, W. Ma, X. Chen, S. Wang and T. Zhang. 2025. MtrAB two-component system is crucial for the intrinsic resistance and virulence of *Mycobacterium abscessus*. Int J Antimicrob Agents 65:107442. 10.1016/j.ijantimicag.2024.107442

27. R. Girase, I. Ahmad and H. Patel. 2024. Bioisosteric modification of Linezolid identified the potential *M. tuberculosis* protein synthesis inhibitors to overcome the myelosuppression and serotonergic toxicity associated with Linezolid in the treatment of the multi-drug resistance tuberculosis (MDR-TB). J Biomol Struct Dyn 42:2111–2126. 10.1080/07391102.2023.2203254

28. W. X. Xu, L. Zhang, J. T. Mai, R. C. Peng, E. Z. Yang, C. Peng and H. H. Wang. 2014. The Wag31 protein interacts with AccA3 and coordinates cell wall lipid permeability and lipophilic drug resistance in *Mycobacterium smegmatis*. Biochem Biophys Res Commun 448:255–260. 10.1016/j.bbrc.2014.04.116

29. P. Plocinski, L. Martinez, K. Sarva, R. Plocinska, M. Madiraju and M. Rajagopalan. 2013. *Mycobacterium tuberculosis* CwsA overproduction modulates cell division and cell wall synthesis. Tuberculosis (Edinb) 93:S21–27. 10.1016/s1472-9792(13)70006-4

30. R. Ravindran, G. Chakrapani, K. Mitra and M. Doble. 2020. Inhibitory activity of traditional plants against *Mycobacterium smegmatis* and their action on Filamenting temperature sensitive mutant Z (FtsZ)-A cell division protein. PLoS One 15:e0232482. 10.1371/journal.pone.0232482

31. I. S. Vadrevu, H. Lofton, K. Sarva, E. Blasczyk, R. Plocinska, J. Chinnaswamy, M. Madiraju and M. Rajagopalan. 2011. ChiZ levels modulate cell division process in mycobacteria. Tuberculosis (Edinb) 91 Suppl 1:S128–135. 10.1016/j.tube.2011.10.022

32. K. C. Rahlwes, S. A. Ha, D. Motooka, J. A. Mayfield, L. R. Baumoel, J. N. Strickland, A. P. Torres-Ocampo, S. Nakamura and Y. S. Morita. 2017. The cell envelope-associated phospholipid-binding protein LmeA is required for mannan polymerization in mycobacteria. J Biol Chem 292:17407–17417. 10.1074/jbc.M117.804377

33. T. Warrier, M. Tropis, J. Werngren, A. Diehl, M. Gengenbacher, B. Schlegel, M. Schade, H. Oschkinat, M. Daffe, S. Hoffner, A. N. Eddine and S. H. Kaufmann. 2012. Antigen 85C inhibition restricts *Mycobacterium tuberculosis* growth through disruption of cord factor biosynthesis. Antimicrob Agents Chemother 56:1735–1743. 10.1128/aac.05742-11

34. F. Wolschendorf, D. Ackart, T. B. Shrestha, L. Hascall-Dove, S. Nolan, G. Lamichhane, Y. Wang, S. H. Bossmann, R. J. Basaraba and M. Niederweis. 2011. Copper resistance is essential for virulence of *Mycobacterium tuberculosis*. Proc Natl Acad Sci U S A 108:1621–1626. 10.1073/pnas.1009261108

35. C. Chhotaray, S. Wang, Y. Tan, A. Ali, M. Shehroz, C. Fang, Y. Liu, Z. Lu, X. Cai, H. M. A. Hameed, M. M. Islam, G. Surineni, S. Tan, J. Liu and T. Zhang. 2020. Comparative analysis of whole-genome and methylome profiles of a smooth and a rough *Mycobacterium abscessus* clinical strain. G3 (Bethesda) 10:13-22. 10.1534/g3.119.400737

36. J. Guo, C. Wang, Y. Han, Z. Liu, T. Wu, Y. Liu, Y. Liu, Y. Tan, X. Cai, Y. Cao, B. Wang, B. Zhang, C. Liu, S. Tan and T. Zhang. 2016. Identification of lysine acetylation in *Mycobacterium abscessus* using LC-MS/MS after immunoprecipitation. J Proteome Res 15:2567–2578. 10.1021/acs.jproteome.6b00116

37. S. Zeng, Y. Ju, M. S. Alam, Z. Lu, H. M. A. Hameed, L. Li, X. Tian, C. Fang, X. Fang, J. Ding, X. Wang, J. Hu, S. Wang and T. Zhang. 2025. A CRISPR-nonhomologous end-joining-based strategy for rapid and efficient gene disruption in *Mycobacterium abscessus*. mLife 4:169–180. 10.1002/mlf2.70007

38. W. Nie, H. Duan, H. Huang, Y. Lu and N. Chu. 2015. Species identification and clarithromycin susceptibility testing of 278 clinical nontuberculosis mycobacteria isolates. Biomed Res Int 2015:506598. 10.1155/2015/506598

39. E. Story-Roller, E. C. Maggioncalda and G. Lamichhane. 2019. Synergistic efficacy of β-lactam combinations against *Mycobacterium abscessus* pulmonary infection in mice. Antimicrob Agents Chemother 63(8):e00614–19. 10.1128/aac.00614-19

