## Supplementary material for "A division-associated envelope protein, MAB_2363, drives intrinsic resistance and virulence in *Mycobacterium abscessus*": Clean Manuscript

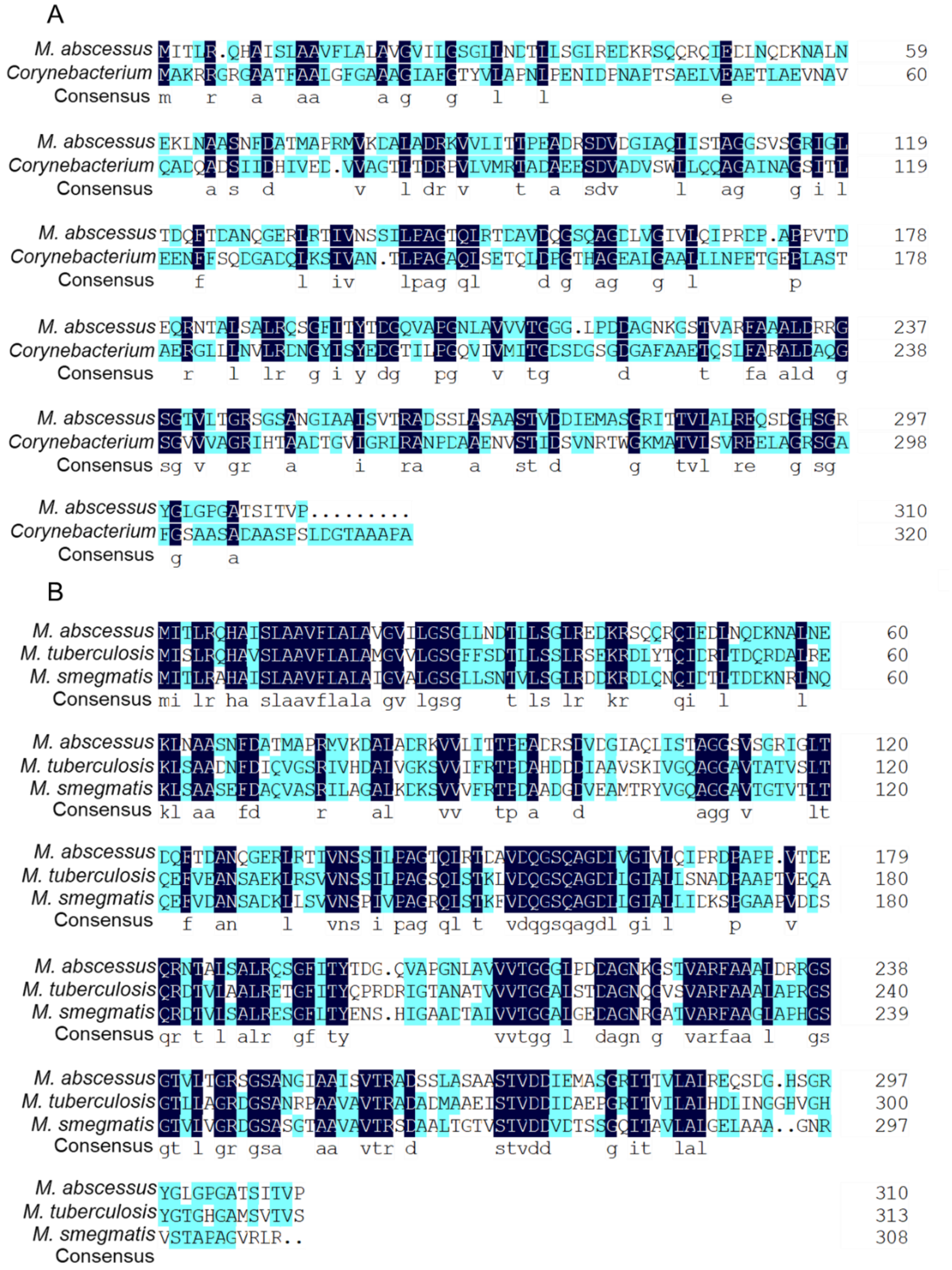

**Figure S1. The alignment of MAB_2363 with SteB and other mycobacterial homologs.** (A) The alignment of MAB_2363 and SteB from *C. glutamicum*. MAB_2363 is 30.98% identical to SteB. (B) The alignment of MAB_2363 with its homologs from other mycobacteria. MAB_2363 is 55.77 and 58.01% identical to its orthologs in *M. tuberculosis* and *M.smegmatis* respectively. Letters highlighted in dark blue color indicate amino acid residues that are identical at that position, light blue signifies residues that differ but share highly similar physicochemical properties, whereas non-highlighted residues are neither identical nor similar.

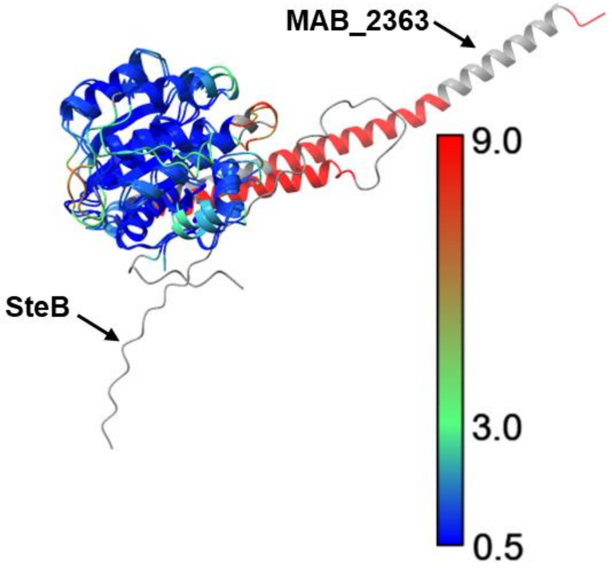


**Figure S2. Structural superposition of core domains from MAB_2363 and *C. glutamicum* SteB.** Superposition of SteB and MAB_2363, with a core root mean square deviation of 0.969 Å (199 atom pairs). The figure was prepared using UCSF ChimeraX 1.10.1.

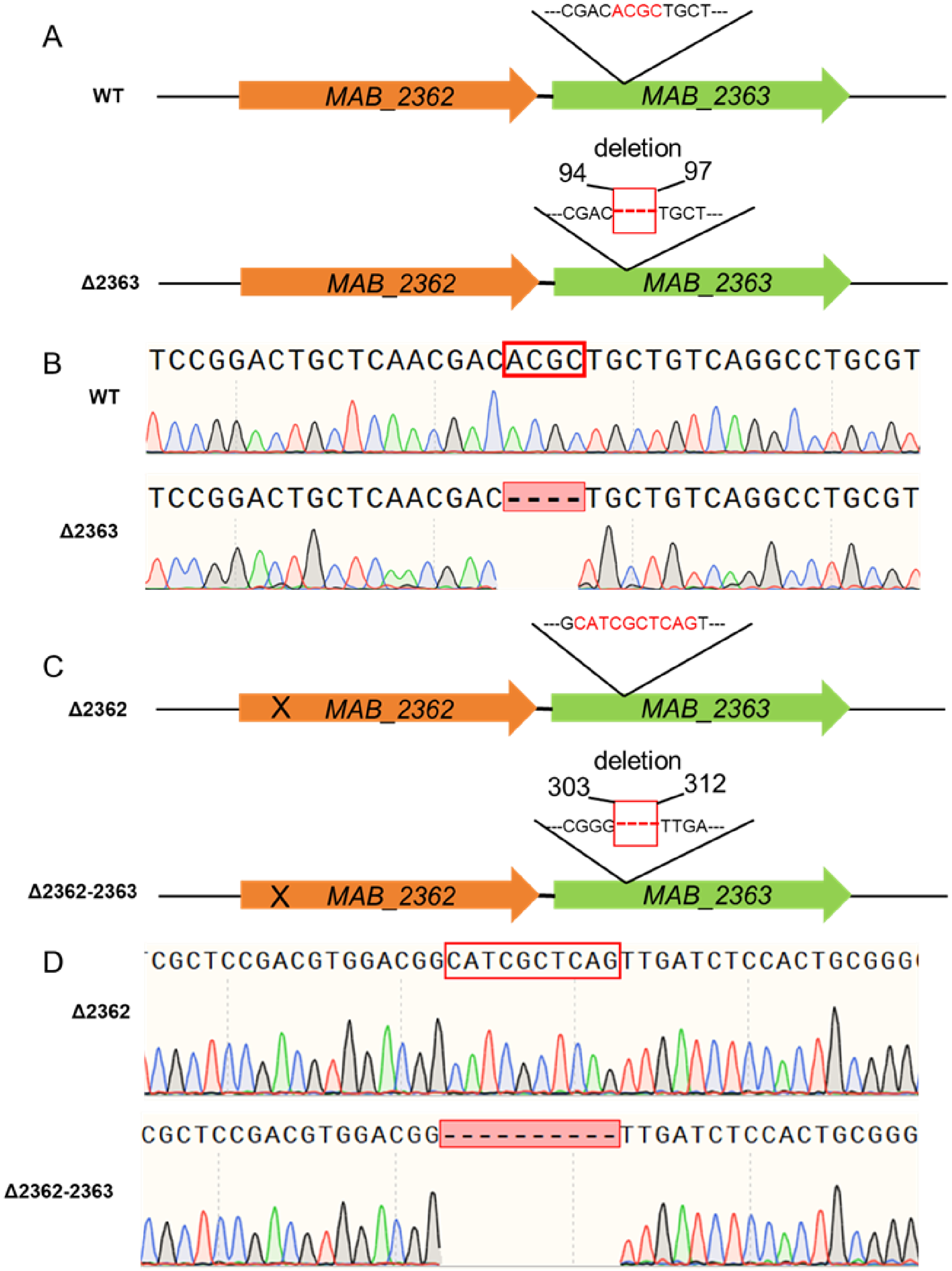


**Figure S3. Gene knockout schematic and sequence alignment plot of *MAB_2363*.** (A) Schematic of the *MAB_2363* gene knockout. The full-length *MAB_2363* is 933 bp, and deletion of bases 94–97 introduced a frameshift mutation, resulting in *MAB_2363* knockout. (B) Sequence alignment plot for *MAB_2363* in the WT and Δ2363 strains. (C) Schematic of the *MAB_2362*-*MAB_2363* double gene knockout. The full-length *MAB_2363* is 933 bp, and deletion of bases 303–312 introduced a frameshift mutation, resulting in *MAB_2363* knockout. (D) Sequence alignment plot for *MAB_2363* in the double knockout strain. WT, wild-type *M. abscessus*; Δ2363, *MAB_2363* knockout strain. Δ2362, *MAB_2362* knockout strain. Δ2362-2363, *MAB_2362-MAB_2363* double knockout strain.

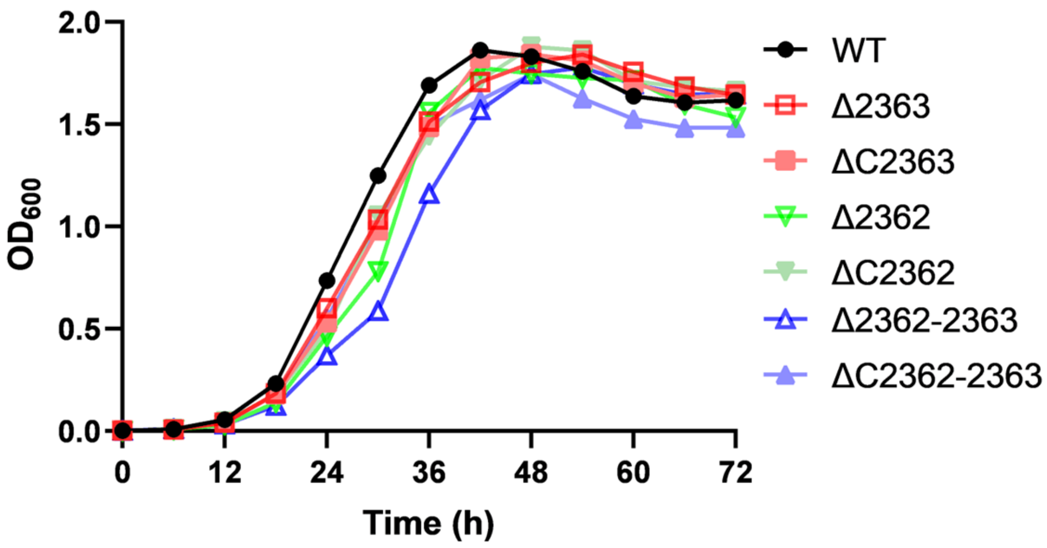


**Figure S4. Growth curves of different *M. abscessus* strains.** WT, wild-type *M. abscessus*; Δ2363, *MAB_2363* knockout strain; ΔC2363, *MAB_2363* complemented strain; Δ2362, *MAB_2362* knockout strain; ΔC2362, *MAB_2362* complemented strain; Δ2362-2363, *MAB_2362-MAB_2363* double knockout strain; ΔC2362-2363, *MAB_2362-MAB_2363* double complemented strain. The experiment was repeated three times.

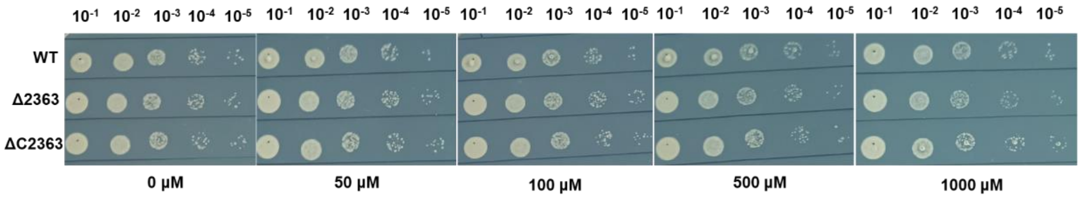


**Figure S5. Susceptibility of different *M. abscessus* strains to $CuSO_4$.** Strains were cultured to the exponential growth phase and streaked onto 7H10 agar plates containing different concentrations of $CuSO_4$. Plates were incubated at 37°C for 3 days before observation. WT, wild-type *M. abscessus*; Δ2363, *MAB_2363* knockout strain; ΔC2363, *MAB_2363* complemented strain.

**Table S1. Cell length of different *M. abscessus* strains.**

| Strains | Cell length (Mean ± SD, μm) |
|---|---|
| WT | 1.85 ± 0.41 |
| Δ2363 | 2.45 ± 0.63 |
| ΔC2363 | 2.17 ± 0.47 |
| Δ2362 | 3.24 ± 0.72 |
| ΔC2362 | 2.08 ± 0.38 |
| Δ2362-2363 | 3.66 ± 0.85 |
| ΔC2362-2363 | 2.30 ± 0.38 |

**Table S2. Bacterial loads in lungs of BDQ- and LZD-treated and untreated mice.**

| Group | $Log_{10}$CFU/Lung (Mean ± SD) | | |
|---|---|---|---|
| | **WT** | **Δ2363** | **ΔC2363** |
| Day 0 | 5.28 ± 0.13 | 5.15 ± 0.12 | 5.12 ± 0.06 |
| Untreated | 5.63 ± 0.17 | 4.52 ± 0.07 | 5.02 ± 0.02 |
| BDQ | 5.63 ± 0.13 | 4.04 ± 0.19 | 4.99 ± 0.16 |
| LZD | 5.66 ± 0.07 | 4.51 ± 0.14 | 5.06 ± 0.06 |

**Table S3. Bacterial burden in the lungs of mice infected with different *M. abscessus* strains.**

| Group | $Log_{10}$CFU/Lung (Mean ± SD) | | | |
|---|---|---|---|---|
| | **Day 0** | **Day 6** | **Day 11** | **Day 16** |
| WT | 5.28 ± 0.13 | 5.64 ± 0.18 | 5.63 ± 0.17 | 6.14 ± 0.10 |
| Δ2363 | 5.15 ± 0.12 | 4.81 ± 0.12 | 4.52 ± 0.07 | 4.10 ± 0.03 |
| Δ2362 | 4.76 ± 0.12 | 3.94 ± 0.17 | 3.21 ± 0.12 | 3.05 ± 0.10 |
| Δ2362-2363 | 4.88 ± 0.09 | 3.87 ± 0.05 | 3.14 ± 0.14 | 2.86 ± 0.10 |

**Table S4. MICs of indicated antibiotics against different *M. abscessus***

**strains.**

| Antibiotics | MIC（μg/mL） | | | | | |
|---|---|---|---|---|---|---|
| | **WT** | **Δ2363** | **ΔC2363** | **ΔCRv1698** | **ΔCMs_3747** | **ΔCV** |
| LZD | 64 | 4 | 64 | 4 | 4 | 4 |
| CLA | 64 | 8 | 64 | 16 | 8 | 8 |
| MXF | 8 | 1 | 4 | 1 | 2 | 2 |
| RFB | 4 | 1 | 2 | 1 | 1 | 1 |
| BDQ | 1 | 0.25 | 0.5 | 0.5 | 0.5 | 0.5 |
| CFX | 32 | 16 | 32 | 16 | 16 | 16 |
| LEV | 32 | 16 | 32 | 16 | 16 | 16 |

WT, wild-type *M. abscessus*; Δ2363, *MAB_2363* knockout strain; ΔC2363, *MAB_2363* complemented strain. ΔCRv1698, *Rv1698* complemented strain; ΔCMs_3747, *MSMEG_3747* complemented strain; ΔCV, empty vector (pMV261) complemented strain. The experiment was performed in triplicates and repeated three times.

**Table S5. Strains used in this study**

| **Strain** | **Description** | **Source** |
|---|---|---|
| WT | *M. abscessus* GZ002 isolated from Guangzhou Chest Hospital | Preserved in our laboratory |
| Mab:pNHEJ-Cpf1 | *M. abscessus* used for constructing gene knockout mutants | Preserved in our laboratory |
| Δ2363 | *MAB_2363* knockout strain | Constructed in this study |
| ΔC2363 | *MAB_2363* complemented strain | Constructed in this study |
| Δ2362-2363 | *MAB_2362-MAB_2363* knockout strain | Constructed in this study |

| | | |
|---|---|---|
| ΔC2362-2363 | *MAB_2362-MAB_2363* complemented strain | Constructed in this study |
| Δ2362 | *MAB_2362* knockout strain | Preserved in our laboratory |
| ΔC2362 | *MAB_2362* complemented strain | Preserved in our laboratory |
| Mab:GFP | *M. abscessus* expressing GFP | Constructed in this study |
| Mab:GFP-MAB_2362 | *M. abscessus* coexpressing GFP and MAB_2362 | Constructed in this study |
| Mab:GFP-MAB_2363 | *M. abscessus* coexpressing GFP and MAB_2363 | Constructed in this study |

**Table S6. Plasmids used in this study**

| **Plasmid** | **Description** | **Source** |
|---|---|---|
| pCR-Zeo | Free *E. coli*—mycobacterium shuttle plasmid containing a temperature-sensitive mycobacterial replicator PAL5000ts, strong promoter *hsp60* driving direct repeats (DR)—target sequence — DR fused with Fn-Cas12a, and ZEO$^{R}$ | Gifted by Laboratory of Yicheng Sun, Peking Union Medical College |
| pCR-Zeo-MAB_2363 | Insertion of *MAB_2363*-targeting crRNA between the two DRs in pCR-Zeo | Constructed in this study |
| pMV261 | Free *E. coli*—mycobacterium shuttle plasmid harboring the strong mycobacterial promoter *hsp60* and KAN$^{R}$ | Preserved in our laboratory |
| pMV261-MAB_2363 | Insertion of *MAB_2363* downstream of *hsp60* in pMV261 | Constructed in this study |
| pMV261-MAB_2362- | Insertion of *MAB_2362-MAB_2363* | Constructed |

| | | |
|---|---|---|
| 2363 | downstream of *hsp60* in pMV261 | in this study |
| pMV261-GFP | Insertion of *gfp* downstream of *hsp60* in pMV261 | Constructed in this study |
| pMV261-GFP-MAB_2362 | Co-insertion of *gfp* and *MAB_2362* downstream of *hsp60* in pMV261 | Constructed in this study |
| pMV261-GFP-MAB_2363 | Co-insertion of *gfp* and *MAB_2363* downstream of *hsp60* in pMV261 | Constructed in this study |

**Table S7. Primers used in this study**

| **Primer** | **Sequence**（5'→3'） | **Annotation** |
|---|---|---|
| cr-MAB_2363-F | CGGACTGCTCAACGACACGCTG | Construction of pCR-Zeo-MAB_2363 |
| cr-MAB_2363-R | CAGCGTGTCGTTGAGCAGTCCG | |
| pMV-yz-F | GAACTCCGTTGTAGTGCTTG | Verification of pMV261-derived plasmids |
| pMV-yz-R | ACGTTTCCCGTTGAATATGG | |
| JD-Zeo-R | GGTGAATCCTCCTGAATATGTAGAG | Verification of pCR-Zeo-MAB_2363 |
| hb-MAB_2362-F | AGACAATTGCGGATCCATGATCACCCCTATGAACTTGACC | Construction of pMV261-MAB_2362-2363 and pMV261-MAB_2363 |
| hb-MAB_2363-F | AGACAATTGCGGATCCAGTGATCACCCTGCGCC | |
| hb-MAB_2363-R | ACTACGTCGACATCGATCTAGGGGACGGTGATGGAGG | |
| pMV-GFP-F | AGACAATTGCGGATCCATGGTGAGCAAGGGCGAG | Construction of pMV261-GFP |
| pMV-GFP-R | ACTACGTCGACATCGATttACTTGTACAGCTCGTCCATGCC | |
| GFP-R | AGGGTGATCGAGCCACCCCCGC | Construction of pMV261-GFP-MAB_2363 |

| | |
|---|---|
| G-MAB_2363-F | TGGCTCGATCACCCTGCGCCAGC |
